# Spatiotemporal dynamics of flow experience: an EEG microstate analysis

**DOI:** 10.64898/2026.05.11.724329

**Authors:** Shiva Khoshnoud, Federico Alvarez Igarzábal, Marc Wittmann

## Abstract

Flow, as defined by Mihalyi Csikszentmihalyi (1975), is a holistic sensation experienced when individuals are fully immersed in an activity, resulting in a mental state characterized by a diminished sense of self and altered perception of time. To investigate the global neural dynamics underlying flow, we employed EEG microstate analysis to capture the spatial and temporal properties of dominant transient global brain states (Lehmann et al., 1998). In a study involving 43 participants playing the video game Thumper for 25 minutes, we extracted three four-minute EEG segments from each session corresponding to reported experiences of flow, boredom, and frustration, as determined by self-reports and performance metrics. Across conditions, six distinct microstate topographies (A–F) accounted for most of the global variance. Given that reduced self-referential processing is a key feature of flow, we hypothesized that flow would modulate the properties of microstates C and E, which have been associated with brain regions resembling the default mode network (DMN). Compared to boredom and frustration, the flow condition showed significantly decreased global explained variance, mean duration, time coverage, and occurrence frequency of microstate E, as well as reduced mean duration and time coverage of microstate C. These findings suggest that microstates associated with self-referential processing are shorter and less frequent during flow than during boredom and frustration. This supports the notion that the flow experience modulates global brain dynamics, particularly within the DMN. Furthermore, our results align with previous research reporting reduced DMN activity during meditative and psychedelic states, reinforcing the idea of diminished self-awareness in such conditions.

## Introduction

There are moments in life when individuals become fully immersed in an activity, experiencing a strong sense of control and deep enjoyment. Csikszentmihalyi (1975, 1990; Csikszentmihalyi et al., 2005) described such instances as “flow.” According to the flow channel model (Csikszentmihalyi, 1975), achieving a balance between an individual’s skill level and the challenge of the activity is essential for the emergence of flow. When the challenge exceeds one’s skills, frustration arises, whereas boredom occurs when skills exceed the level of challenge. Experiencing flow is considered integral to human well-being, as it enriches individuals’ lives (Csikszentmihalyi et al., 2005). In the present study, we examine these three states—flow, boredom, and frustration—and compare their neural correlates using EEG microstate analysis.

Despite the significance of flow in everyday life, research on its underlying neurophysiological mechanisms has yielded heterogeneous and sometimes inconsistent findings (for a review, see Khoshnoud et al., 2020). The flow state is generally described as a positive mental state characterized by heightened arousal (Chanel et al., 2011; De Sampaio Barros et al., 2018; Keller et al., 2011; Tian et al., 2017), focused attention (Harris et al., 2016; Klasen et al., 2012; Ullén et al., 2010; Ulrich et al., 2014, 2016; Yoshida et al., 2014), and synchronized activity within the brain’s attention and reward networks (Castellar et al., 2016; De Sampaio Barros et al., 2018; Huskey et al., 2018; Ju & Wallraven, 2019; Klasen et al., 2012; Ulrich et al., 2014, 2016; Yoshida et al., 2014). This state is associated with more automatic action control and reduced self-referential processing (De Sampaio Barros et al., 2018; Ju & Wallraven, 2019; Sadlo, 2016; Ulrich et al., 2014, 2016, 2018). A key neural correlate of this experience is the downregulation of the default mode network (DMN), particularly the medial prefrontal cortex (mPFC), which is closely linked to self-referential processing (Khoshnoud et al., 2020). The DMN is widely considered a central hub associated with the narrative (self-referential) self (Northoff et al., 2006).

In addition to the possibility that these divergent findings can be explained by distinct subprocesses of flow (e.g., attentional focus, reward processing, reduced self-awareness), inconsistencies in previous studies may also stem from the rapid temporal dynamics of large-scale neural networks associated with flow. Such dynamics may not be adequately captured by the low temporal resolution of functional magnetic resonance imaging (fMRI) or by the predominantly local measures typically derived from electroencephalography (EEG).

Although the global field power of EEG provides a measure of overall brain activity, commonly used local metrics—such as amplitude and spectral power—exhibit substantial moment-to-moment variability, which can complicate the interpretation of localized differences during flow. In contrast, EEG microstates represent global patterns of scalp potential topography that remain stable for brief periods (approximately 80–120 ms) before transitioning to a different configuration (Lehmann et al., 1987; Michel & Koenig, 2018). Described by Lehmann and colleagues (1998) as the “building blocks of spontaneous thinking” or “atoms of thought,” and conceptualized as functional moments of temporal integration (Wittmann, 2011), these transiently stable states have proven valuable across a wide range of research applications.

Researchers have employed EEG microstates and their temporal characteristics to investigate the content of spontaneous thoughts (Bréchet et al., 2019; Lehmann et al., 1998, 2010), characterize specific cognitive processes (Milz et al., 2016; Seitzman et al., 2017), identify different sleep stages (Brodbeck et al., 2012), and detect alterations associated with neuropsychiatric disorders (D’Croz-Baron et al., 2019; Nishida et al., 2013; Rieger et al., 2016; Stevens et al., 1997). Furthermore, EEG microstates have proven valuable in studying altered states of consciousness, including meditation and hypnosis (Bréchet & Michel, 2022; Faber et al., 2017; Katayama et al., 2007; Panda et al., 2016). Among the established microstate classes, microstates E and C have been associated with brain regions commonly attributed to default mode network (DMN) activity (Custo et al., 2017; Michel & Koenig, 2018; Panda et al., 2016).

The phenomenon of flow involves parallel processing across distributed brain networks, including attentional networks and the default mode network (DMN), among others (Khoshnoud et al., 2020). This complexity necessitates approaches that capture the dynamics of whole-brain neural activity. In the present study, we introduce EEG microstate analysis as a promising tool for investigating the multimodal characteristics of the flow experience.

Using a sample of 43 participants engaged in an ecologically valid paradigm (video game play), we examined the dynamic properties of EEG microstate maps during periods corresponding to experiences of flow, boredom, and frustration. We hypothesized that key features of the flow state—such as reduced self-referential processing and heightened attentional focus—would modulate the temporal and spatial dynamics of microstate maps, particularly those associated with activity in DMN-related regions.

## Materials and Methods

### Participants and Experimental design

The EEG dataset used in this study comprises 43 recordings obtained from 22 recreational gamers and 21 non-gamers while they played the commercial video game Thumper (for further details on the game and its dynamics, see the behavioral analysis in Rutrecht et al., 2021, and the standard neurophysiological analyses in Khoshnoud et al., 2022). Thumper is a fast-paced rhythm game in which players synchronize button presses with a musical soundtrack and visual stimuli. Players control a silver beetle that moves automatically along a track and must respond to obstacles encountered along the way.

Participants completed two consecutive 60-minute training sessions on separate days. In a third and final session, they played the game for 25 minutes while EEG and cardiorespiratory signals were recorded. The study was approved by the local Ethics Committee of the Institute for Frontier Areas of Psychology and Mental Health (IGPP-2019-01), and all participants provided informed consent prior to participation.

### Retrospective evaluation of the experience

To delineate time intervals associated with the experiences of boredom, flow, and frustration during continuous gameplay, we assessed participants’ experiences using two quantitative measures. The first measure consisted of minute-by-minute self-reports of flow and perceived balance between skill and challenge. Following gameplay, participants reviewed a recording of their performance and provided retrospective minute-by-minute ratings of perceived challenge relative to their skill level using a nine-point Likert scale ranging from −4 (too little challenge) to +4 (too much challenge). Flow experience was similarly rated on a nine-point Likert scale ranging from 1 (not at all) to 9 (completely).

The second measure involved an experimenter-based evaluation of participants’ performance on a minute-by-minute basis. This evaluation included recording the number of errors, instances of damage, and “death” events (i.e., crashes) occurring in each minute of gameplay. Based on the combination of self-reported measures and performance evaluations, we identified and extracted three four-minute segments per participant from both the behavioral data and the EEG recordings, corresponding to flow, boredom, and frustration conditions.

A flow segment was defined as the interval with the highest flow rating, a balanced challenge-to-skill ratio (subjectively reported), and the lowest number of errors (objectively measured). In contrast, boredom segments were characterized by the lowest flow ratings, minimal errors, and a perceived challenge level below the participant’s skill. Frustration segments were defined by a higher number of errors, mid-to-low flow ratings, and a perceived challenge level exceeding the participant’s skill.

### Signal recording and preprocessing

We collected EEG data using a 32-channel electrocap (ActiCHamp, BrainVision), with electrodes positioned according to the extended 10-10 international system (American Electroencephalographic Society, 1993). The Fz electrode served as the reference, and electrode impedances were maintained below 10KΩ. Simultaneous EEG recording was conducted at a sampling rate of 1000 Hz, with online filtering applied within the 0.01 - 120 Hz range.

The preprocessing of EEG data is carried out using a combination of custom-written scripts in MATLAB (The MathWorks, R2019a) and the EEGLAB toolbox (Delorme & Makeig, 2004). Initially, the recorded EEG signals were down-sampled to 250 Hz and filtered using a band pass filter of 1 - 40Hz. Subsequently an artifact subspace reconstruction approach (ASR; Kothe & Jung, 2014) removed transient and large amplitude artifacts. ASR is a principle component-based artifact removal method that identifies clean portions to stablish thresholds for rejecting components (Chang et al., 2020). After these steps, EEG signals underwent independent component decomposition using adaptive mixture independent component analysis (AMICA; Palmer et al., 2008, 2011). The equivalent current dipoles of each independent component (IC) were computed using the three-shell, boundary-element-method head model based on the MNI brain template employing the DIPFIT plugin of the EEGLAB toolbox. Automatic classification and labeling of ICs were performed using the ICLabel plugin of the EEGLAB toolbox employing a machine-learning approach to classify the ICs based on spectral properties and brain topography (Pion-Tonachini et al., 2019). The information derived from the IC labels together with the dipole residual variance was then used to preserve pure brain-related activity while eliminating those components that are associated with eye-blinks, muscle noise, heart artifacts, and other contamination.

### Microstate analysis

Microstate analysis is performed using the free academic software Cartool (Brunet et al., 2011). The general approach for determining microstate maps, which is shown in Figure 1, comprises of two steps. First, the segmentation step in which the most dominant topographies of brain electrical activity across all individuals within a condition was calculated by means of a clustering approach. In the second stage, these dominant microstate maps are fitted back to the EEG data by assigning each data point of the original EEG to a dominant map based on the spatial correlation between the corresponding map and each data point.

**Figure 1.**
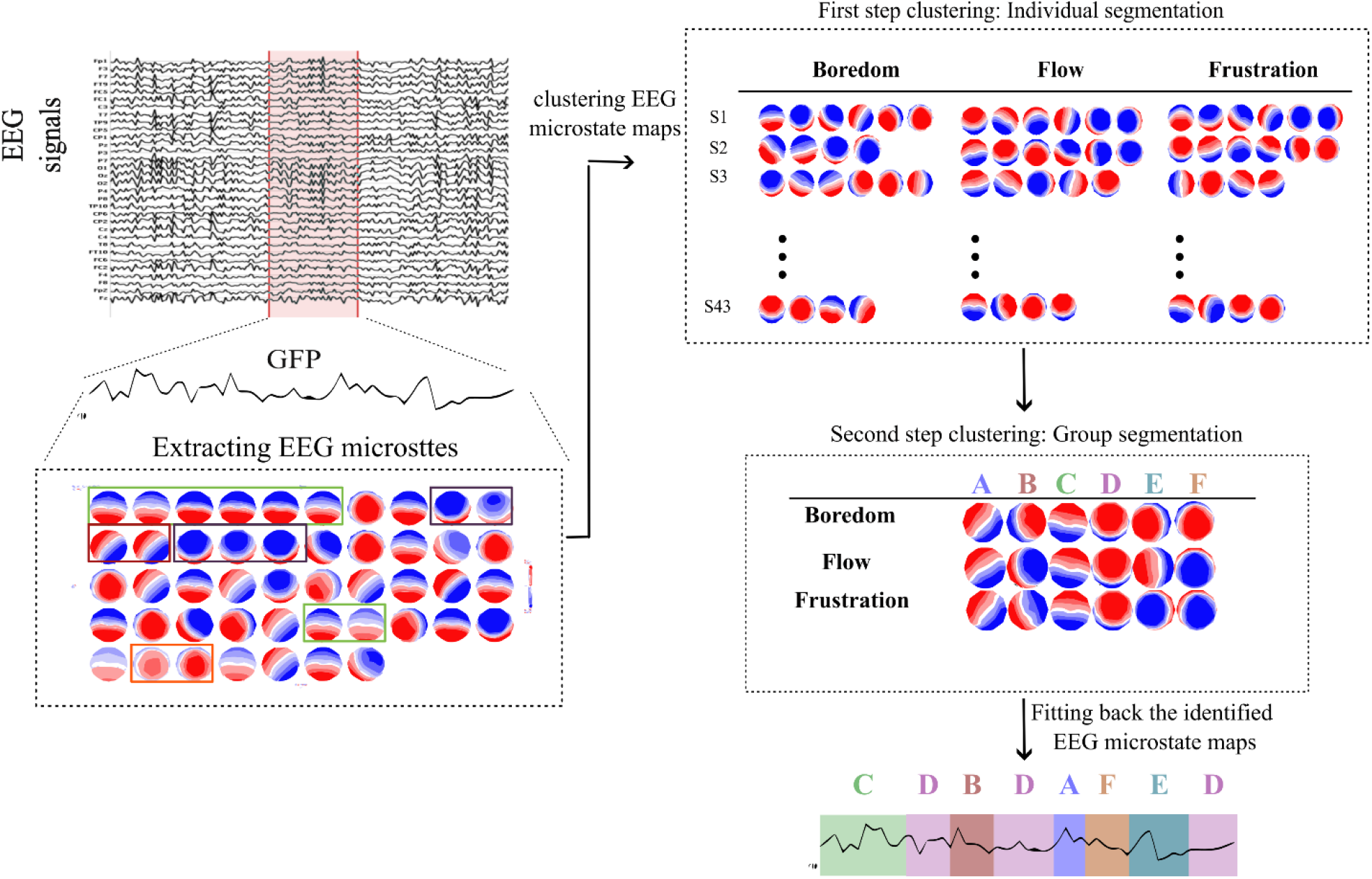
Schematic illustration of the EEG microstate analysis. See text for details.

We calculated the global filed power (GFP) of the EEG data at each time point (*u* = {*u*_1_, *u*_2_, …, *u*_*N*_} with N = 32 scalp electrodes) according to Equation 1 and extracted the topographical maps of the brain electrical field at local maxima of the GFP values for each participant. The GFP peaks were chosen as they represent moments of highest topographical signal-to-noise ratio (Lehmann et al., 1998).

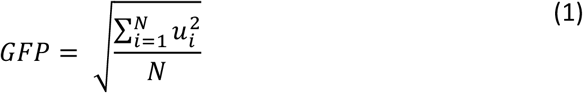

The extracted topographies were then submitted to a modified k-means cluster analysis (Pascual-Marqui et al., 1995), ignoring polarity inversion, to identify the most dominant microstate maps. The clustering was performed in two steps: first on the data of each individual and each condition separately and then on the topographical maps of individuals within a condition. This way the most dominant maps within each condition were calculated (see Figure 1).

One essential factor is determining the optimal number of clusters. We selected this number based on a meta-criterion parameter available in the CARTOOL software (Custo et al., 2017). This index encompasses seven different criteria taken from the literature such as the cross-validation criterion (Pascual-Marqui et al., 1995) or Krznowski-Lai index (Brunet et al., 2011).

To fit back the group microstate maps to the corresponding original EEG time series, the spatial correlation between the group maps of each condition and the extracted topography in each time point was calculated and the time point with the highest correlation was labeled with the map (Michel & Koenig, 2018). If the spatial correlation of a time point with all maps was less than 0.5, then no label was assigned. We applied temporal smoothing by rejecting small segments where the assigned microstate map was present less than 4 time frames (15.6 ms). Once the labels were assigned, we calculated the following parameters for each individual in each condition:

1. GEV – Global Explained Variance; EV= Explained Variance (the squared spatial correlations between a given microstate map and the topography of the corresponding labeled data points) weighted by the GFP (Global Field Power) at each time point as follows:

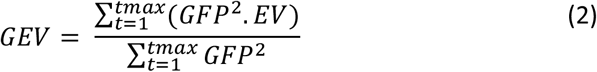 This measure reflects how well a microstate map can describe the data points.
2. Mean microstate duration – The average duration, in milliseconds, that a microstate map is continuously present. This index provides insights into the temporal stability of a microstate map. Longer mean durations might suggest a more stable or sustained map, while shorter mean durations of a microstate might indicate a more rapidly changing brain state.
3. Time coverage of microstate – The proportion of time that a microstate is present during the whole length of corresponding EEG data. Time coverage reflects the dominance of a certain microstate over other microstates during the recording. Larger time coverage can indicate a more significant role of that EEG map on the brain activity during the recording.
4. Mean frequency of occurrence – The average frequency that a microstate is present. Mean frequency of occurrence offers a quantitative measure of how often the brain’s electrical activity patterns transition into specific states.
5. EEG microstate syntax – This measure refers to the sequential transition between EEG microstates and is calculated as transition probabilities from the preceding state to the current state for each microstate pair. These transition probabilities can indicate the brain’s preferred states or sequences during an activity or recording. This study focused on the transition probability of all maps to map E and from map E to other maps, as microstate E has been associated with the regions in the default mode network (DMN). Therefore, we extracted and examined the probabilities of transitioning from maps A, B, C, D, and F to map E and vice versa for further analysis.

### Statistical analysis

One-way repeated measure ANOVAs were used to compare the temporal parameters of each microstate map between three conditions. Degrees of freedom were corrected with the Greenhouse-Geisser correction method. For each parameter and each map, we conducted a repeated measure ANOVA with a threshold of p<0.001. Whenever the ANOVA results were significant, the pair-wise comparisons were calculated using Bonferroni correction.

To statistically assess the transition probabilities of extracted microstate pairs (all maps to E and from E to all other maps) between three conditions, we used a 3*5 repeated measure ANOVA with condition (Boredom, Flow, and Frustration) and microstate pair as two within-subject factors. Whenever the ANOVA results were significant, the pair-wise comparisons were calculated using Bonferroni correction.

## Results

After extracting microstate topographies at local maxima of the GFP for each individual, we applied the modified k-means clustering analysis first to each individual EEG data and then across subjects within a condition. While the first step led to different optimal numbers of maps across individuals (4 to 7 maps), the second clustering analysis within each condition using the meta-criterion parameter identified 6 optimal microstate maps for each condition (see Figure 1). These 6 topographies explained 65-70% of the global variance. As one can see from Figure 1, the microstate maps of the three conditions of boredom, flow, and frustration are strikingly similar to each other and correspond well to the established six microstate maps (A-F) in previous studies (Bréchet et al., 2019; Michel & Koenig, 2018). After getting the most dominant microstate maps of each condition, we fitted back group maps of each condition to the corresponding EEG data separately and then we calculated the 4 parameters of GEV, mean duration, time coverage, and mean frequency of occurrence for each map.

### Global Explained Variance (GEV)

Regarding the GEV, a one-way repeated-measure ANOVA revealed a significant main effect of condition for both microstate maps E and F (F (1.14,47.9) =26.41, p <0.001, η2= 0.38 ; F(1.55,65.4) =23.38, p <0.001, η2= 0.35, respectively). As shown in Figure 2a and 2b, the GEV of map E decreased during the flow condition compared to the boredom (p=0.004) and the frustration (p<0.001) conditions. However, the GEV of map F increased during the flow condition compared to the boredom (p=0.02) and the frustration (p<0.001) conditions.

**Figure2.**
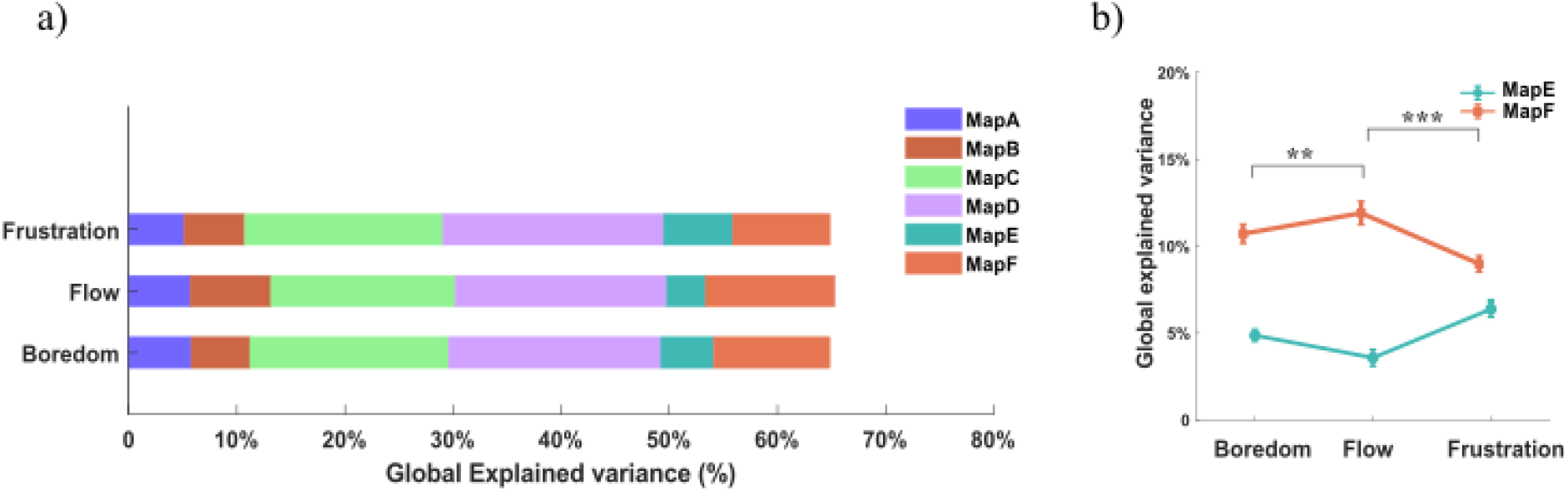
a) The Global explained variance (GEV) of the six dominant microstate maps extracted for three conditions of boredom, flow, and frustration. b) The average GEV of microstate maps E and F were significantly modulated during flow compared to boredom and frustration.

**Figure3.**
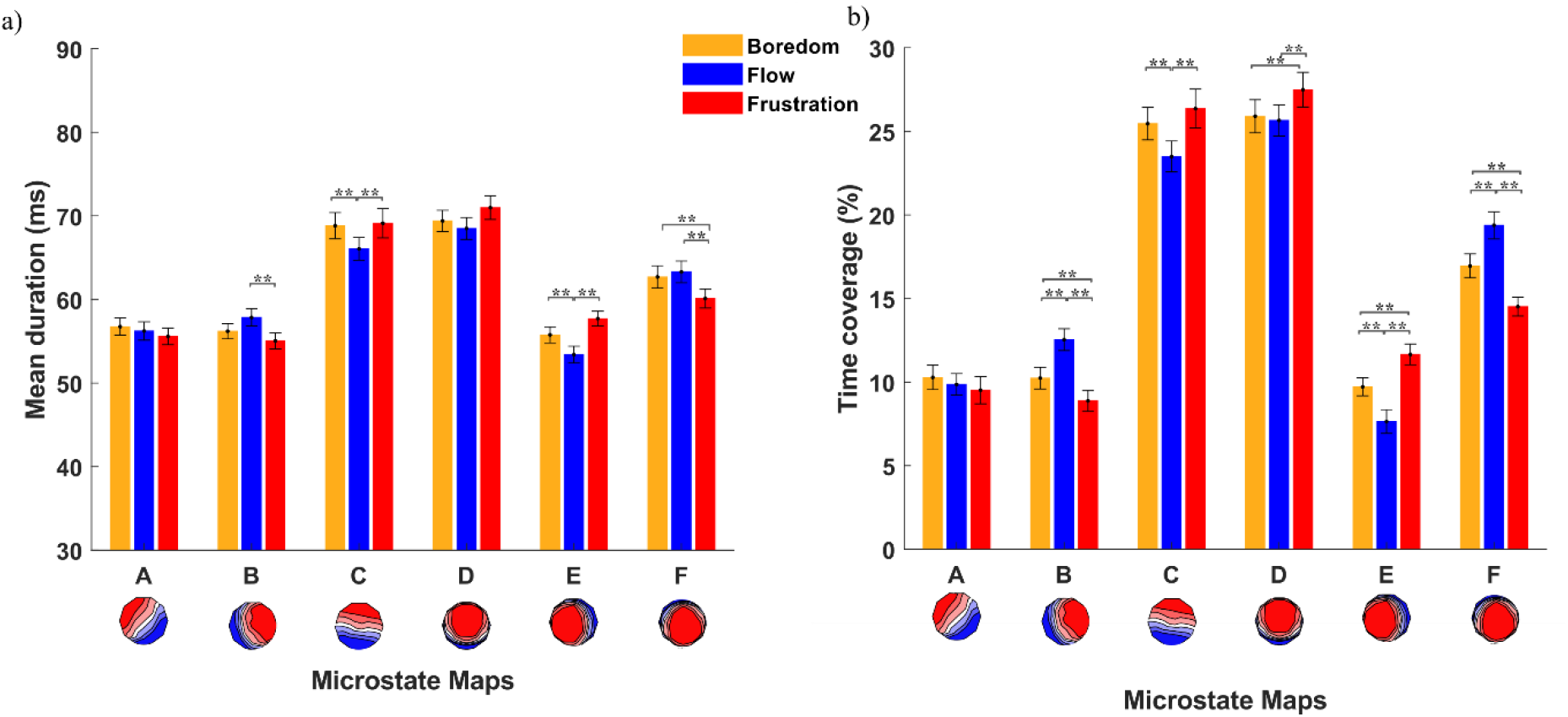
a) Mean and standard error of the average duration of each microstate map during three conditions of boredom, flow, and frustration. b) Mean and standard error of the average percentage of time coverage for each microstate map during three conditions.

**Figure4.**
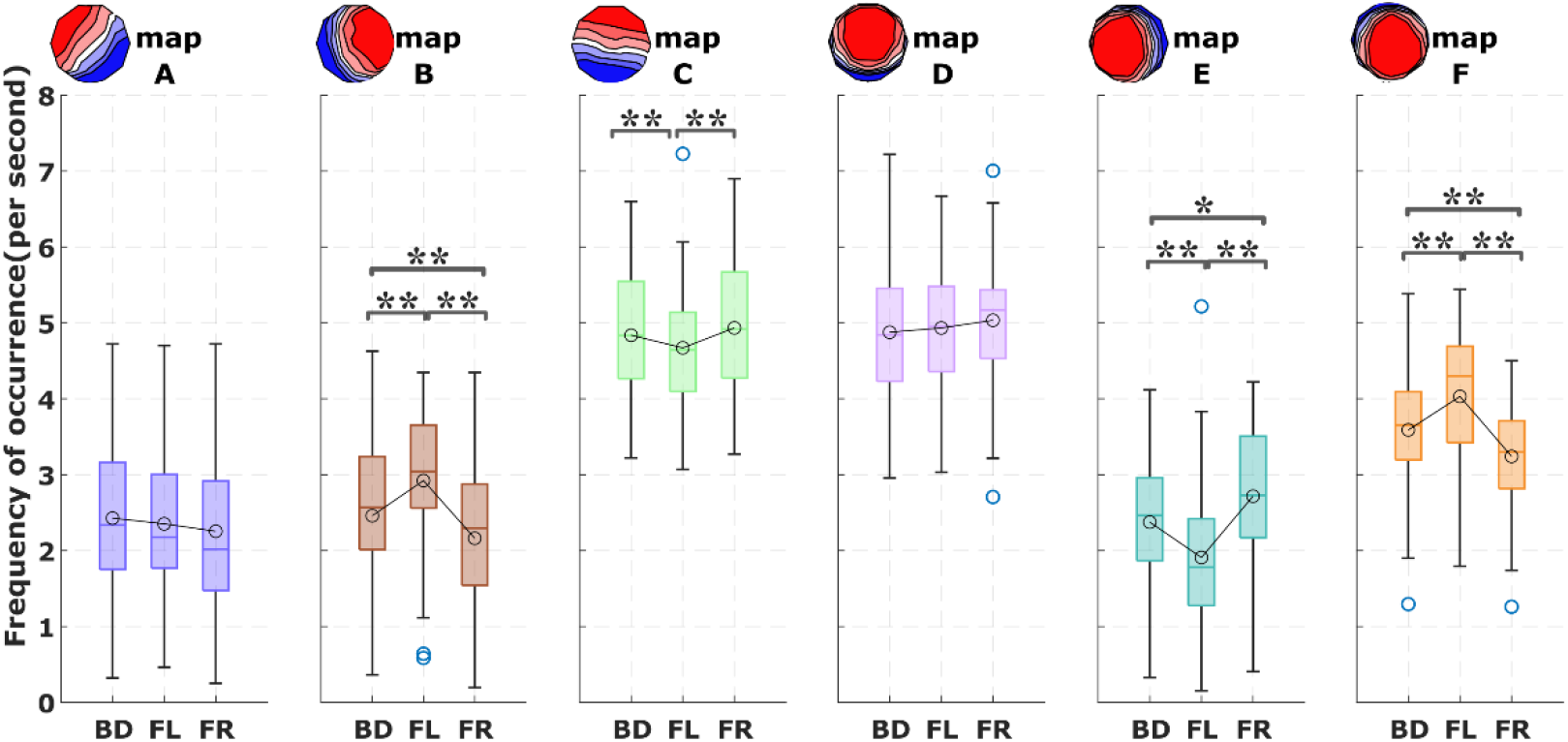
Boxplots of frequency of occurance for each microstate map during three conditions of boredom, flow, and frustration.

### Mean microstate duration

Regarding mean duration of each microstate, we found a significant main effect of condition (boredom, flow, frustration) for the microstate maps B, C, E, and F (F(2,84) =7.92, p <0.001, η2= 0.15 ; F(2,84) =9.36, p <0.001, η2= 0.18; F(2,84) =21.31, p <0.001, η2= 0.33 ; F(1.75,73.3) =11.64, p <0.001, η2= 0.21, respectively). Bonferroni corrected post-hoc tests showed that microstate B increased in mean duration in the flow condition compared to the boredom (p=0.06) and to the frustration (p<0.001) conditions. The mean duration of microstates C and E significantly decreased during flow compared to the boredom (p=0.002, p=0.002) and frustration (p<0.001, p<0.001) conditions. For microstate F, we observed a significant decrease in the mean duration during frustration compared to flow (p<0.001) and boredom (p=0.001).

### Time coverage of microstates

We observed a main effect of condition for the time coverage of microstates B, C, D, E, and F (F(1.15,48.48) =44.85, p <0.001, η2= 0.51 ; F(1.57,66.06) =20.36, p <0.001, η2= 0.32; F(2,84) =15.86, p <0.001, η2= 0.27 ; F(1.13,47.67) =32.27, p <0.001, η2= 0.43; and F(1.61,67.82) =47.53, p <0.001, η2= 0.53,respectively). Time coverage of microstates B and F significantly increased during flow compared to the boredom (both p<0.001) and the frustration (both p<0.001) conditions. In both maps (B, F), time coverage during the frustration was significantly lower compared to the boredom condition (both p<0.003).

For the microstates C and E, post-hoc tests revealed that time coverage of these microstates was significantly decreased during the flow condition compared to the boredom (both p<0.001) and the frustration (both p<0.001) conditions. Time coverage of microstate E was significantly higher during the frustration condition compared to the boredom (p<0.001). Finally, for microstate D, there was higher time coverage during frustration condition compared to the flow (p<0.001) and boredom (p<0.001) conditions.

### Mean frequency of occurrence of microstates

When comparing the frequency of occurrence, the most prominent finding was the significantly distinct occurrence rate for microstates B and F (F(1.23,51.79) =51.37, p <0.001, η2= 0.55 ; F(1.63,68.73) =52.9, p <0.001, η2= 0.55) with increased occurrence rate of both microstates during the flow compared to the boredom (both p<0.001) and the frustration (both p<0.001) conditions. Frequency of occurrence for these microstates was significantly lower during the frustration in comparison to the boredom condition (both p<0.0001). Microstate E significantly decreased in occurrence during the flow (F (1.17,49.40)=31.26, p <0.001, η2= 0.42) compared to the boredom and the frustration conditions (both p<0.001). For microstate E the occurrence rate was significantly higher in the frustration condition than for boredom (p=0.004). There was also a distinct occurrence rate for the microstate C (F (2,84) =8.29, p <0.001, η2= 0.16) with a lower rate in the flow condition compared to the frustration (p<0.001) and boredom (p=0.038) conditions.

### Syntax of EEG microstates

In Figure 5a and 5d, the average transition probabilities, along with standard deviations, are depicted for the transitions between five microstates and microstate E, during the boredom, flow, and frustration conditions. The application of a repeated measure ANOVA on observed transition probabilities from all other microstates to E revealed a significant main effect of condition (F (1.11,46.85) = 43.62, p<0.001, η^2^= 0.36). This indicates overall lower transition probabilities to state E during flow compared to frustration (p<0.001) and boredom (p<0.001), as illustrated in Figure 5a. Frustration also resulted in higher transition probabilities from other microstates to E compared to boredom (p<0.001).

**Figure5.**
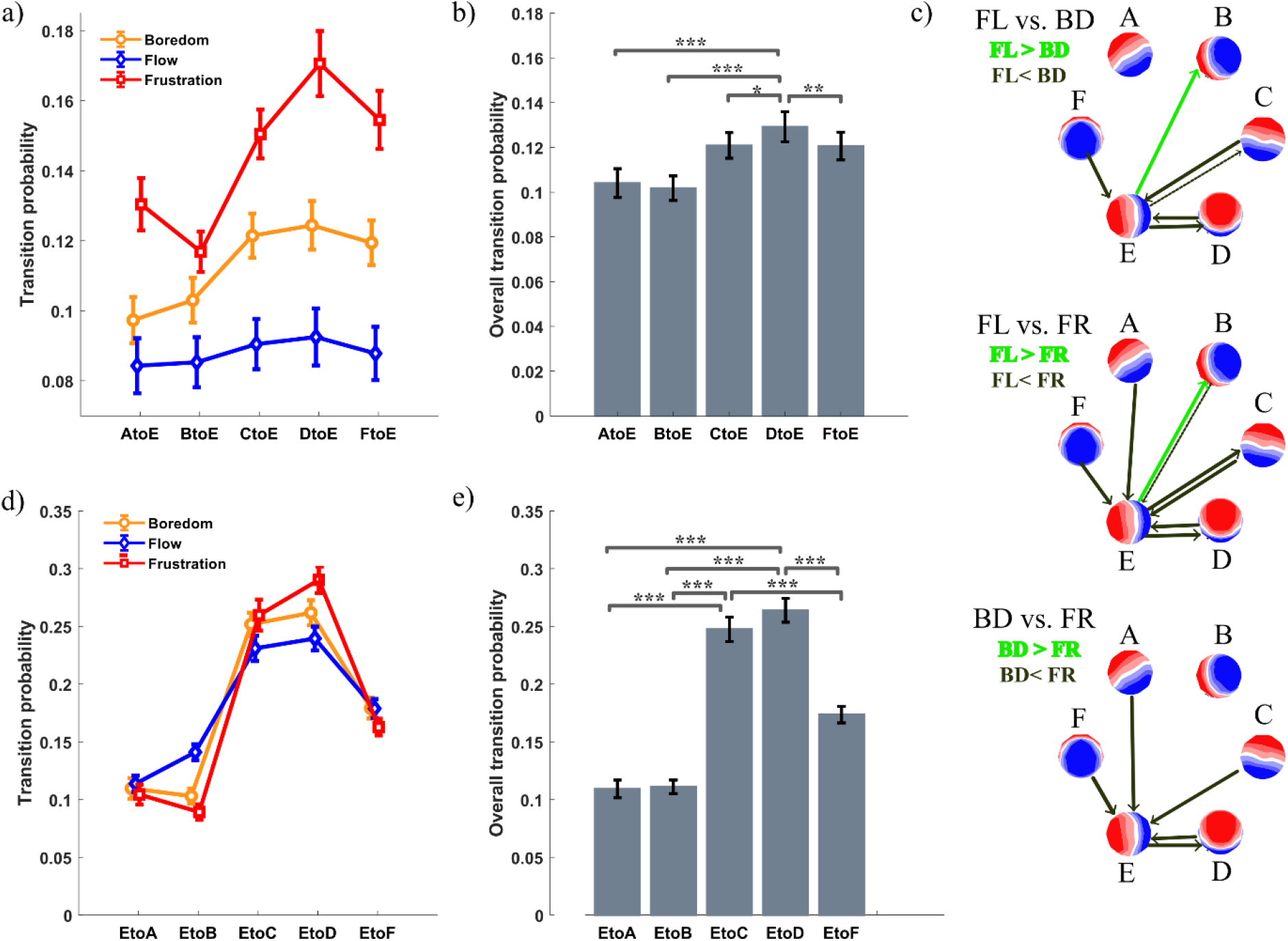
a) Average and standard error of observed transition probabilities from five microstates (A, B, C, D, and F) to microstate E during the three conditions of boredom, flow, and frustration; b) overall transition probabilities regardless of condition from other states to microstate E; c) comparison of the transition probabilities from other states to E between three conditions; d) average and standard error of observed transition probabilities from microstate E to other five microstates (A, B, C, D, and F during the three conditions; e) overall transition probabilities regardless of condition from the microstate E to other states.

Additionally, a significant main effect of microstate pair was observed (F (4,168) =40.59, p<0.001, η2= 0.07), emphasizing that microstate E was significantly more often followed by microstate D compared to others (all p<0.01; see Figure 5b). Post hoc analysis revealed that transitions from C, D, and F to E were significantly lower during flow compared to boredom and frustration (see Figure 5c).

A similar analysis was conducted to compare transition probabilities from microstate E to the other five microstates between the conditions. Figure 5d shows that the probabilities of microstate E transitions to other five microstates did not exhibit a significant difference between conditions (F (2,84) = 0.18, p=0.83, η^2^< 0.01). Notably, a strong main effect of microstate pair was observed (F (2.72,114.3) = 61.02, p<0.001, η^2^= 0.53), indicating that microstate E transitioned significantly more frequently to microstates C, and D compared to any other state (all p<0.001). However, microstate E showed lower transition probabilities to states C and D during flow compared to boredom and frustration. This microstate was more often followed by microstate B during Flow compared to boredom and frustration (p<0.001).

## Discussion

By investigating EEG microstates during a gameplay session, this study offers compelling evidence of altered spatiotemporal dynamics in large-scale brain networks during the experience of flow, boredom, and frustration. In a 25-minute-long gameplay (playing the video game Thumper) we recorded EEG signals of 43 participants and based on retrospective self-reported measures and performance evaluations, we extracted EEG segments corresponding to the experiences of flow, boredom, and frustration for each individual. The observed dominant EEG microstate maps across these states showed substantial similarity and aligned with those described in prior research (Bréchet et al., 2019; Michel & Koenig, 2018).

Building on the hypothesis that flow is characterized by reduced self-referential processing (Csikszentmihalyi, 1990; Khoshnoud et al., 2020), we anticipated changes in the temporal and spatial dynamics of microstate maps C and E, which are linked to default mode network (DMN) activity (Custo et al., 2017; Panda et al., 2016). Our results support this hypothesis, demonstrating that microstates C and E exhibited reduced temporal stability (as indicated by shorter mean duration), diminished dominance (as reflected in lower time coverage), and decreased occurrence (as shown by frequency of occurrence) during flow compared to the boredom and frustration.

Previous research has demonstrated variability in the duration of microstate C under different states of consciousness. Specially, a decreased duration of microstate C has been observed during light hypnosis (Katayama et. al. 2007) and under light surgical anesthesia (Artoni et al., 2022). In contrast, an increased duration and occurrence of microstate C have been documented in scenarios such as deep anesthesia (Artoni et al., 2022), during tasks involving autobiographical memory (Bréchet et al., 2019), while dreaming during NREM sleep (Bréchet et al., 2020), and throughout meditation states (Panda et al., 2016). All of these scenarios involve self-consciousness and self-referential processing.

Very few studies have investigated microstate E (Bréchet et al., 2019; Custo et al., 2017) and its associated network, which includes regions such as the anterior cingulate cortex (ACC) and medial prefrontal cortex (mPFC). In a recent systematic review, Tarailis et al. (2024) suggested that microstate E (referred to as microstate F in their nomenclature) is linked to the anterior default mode network (DMN) and plays a role in processing personally relevant information. The observed reduction in the temporal characteristics of microstate E during flow in the present study may therefore reflect a decreased engagement with self-relevant and internally directed cognition, consistent with the phenomenological hallmark of diminished self-awareness during flow states.

In addition to the reductions observed in microstates C and E, we found increased temporal dominance (i.e., duration, time coverage, and occurrence) of microstates B and F during flow. Previous studies have associated microstate B with visual processing networks and externally oriented attention, suggesting enhanced sensory engagement with the task at hand, in our study referring to playing the game *Thumper*. Similarly, microstate F has been linked to attentional and salience-related processes, potentially reflecting increased integration of task-relevant information. Together, these findings suggest a shift from internally oriented processing toward externally focused, task-relevant neural dynamics during flow.

The observed alterations in microstate syntax further support this interpretation. Specifically, the reduced transition probabilities toward microstate E during flow indicate a decreased likelihood of the brain entering DMN-related states. This reduced accessibility of self-referential processing states may contribute to the sustained attentional engagement and action automatization characteristic of flow. Conversely, the increased transitions involving task-relevant microstates (e.g., B and F) suggest a more stable and efficient neural configuration supporting ongoing performance.

These findings align with a previous neuroimaging review reporting decreased DMN activity and increased engagement of attentional and task-positive networks during flow (Khoshnoud et al., 2020). Moreover, the present results are consistent with research on other altered states of consciousness, such as meditation and psychedelic experiences, which have also been associated with reduced DMN activity and diminished self-referential processing (Wittmann, 2022; 2025). This convergence across different domains supports the notion that reduced dominance of DMN-related microstates may represent a general neural signature of states characterized by attenuated self-awareness, a key feature of flow states (Csikszentmihalyi, 1975, 1990; Csikszentmihalyi et al., 2005).

Importantly, the use of EEG microstate analysis in this study provides novel insights into the fast temporal dynamics underlying the flow experience. Unlike fMRI, which is limited in temporal resolution, microstate analysis captures rapid transitions between global brain states on the millisecond scale. This allows for a more fine-grained characterization of the dynamic interplay between large-scale neural networks during complex experiential states such as flow.

Several limitations of the present study should be acknowledged. First, the identification of flow, boredom, and frustration segments relied partly on retrospective self-reports, which may be subject to recall biases. Although combined with objective performance measures, future studies could benefit from real-time assessments of subjective experience. In one recent study employed a novel method, continuous and real-time subjective measurement of flow was conducted using a pedal that indicated felt flow during playing the video game *Thumper* (Genc et al., 2026): Results supported the hypotheses that interacting with the pedal did not interfere with players’ flow experience as indicated by the correlation between real-time and post-task flow ratings. Second, the use of a specific video game paradigm, while ecologically valid, may limit the generalizability of the findings to other types of flow-inducing activities.

Future research could extend these findings by combining EEG microstate analysis with other neuroimaging techniques, such as fMRI, to better link temporal dynamics with spatially resolved brain networks. Additionally, investigating individual differences in flow propensity and expertise (e.g., gamers vs. non-gamers) may further elucidate how neural dynamics support optimal experience. Finally, exploring interventions that modulate microstate dynamics may provide novel avenues for enhancing flow in applied settings, such as education, sports, and clinical contexts.

In conclusion, the present study demonstrates that the flow experience is characterized by distinct alterations in the temporal dynamics of EEG microstates, particularly those associated with the default mode network. Reduced dominance and accessibility of DMN-related microstates, alongside increased engagement of task-relevant states, support the notion that flow involves a shift from self-referential to externally oriented processing. These findings highlight the value of EEG microstate analysis as a powerful tool for investigating the dynamic neural basis of complex experiential states.

## Author Contributions

All authors contributed to conceptualization, design of the study, manuscript revision, and all approved the submitted version. SK performed data curation, conducted the analysis, and wrote the first draft.

